# The enzyme glutamate-cysteine ligase (GCL) is a target for ferroptosis induction in cancer

**DOI:** 10.1101/2024.04.28.591552

**Authors:** John K. Eaton, Priya Chatterji, Laura Furst, Sneha Basak, Ayesha M. Patel, Yan Y. Sweat, Luke L. Cai, Krishna Dave, Rachelle A. Victorio, Elizabeth Pizzi, Javad Noorbakhsh, Gaochao Tian, Jennifer A. Roth, John Hynes, Gang Xing, Mathias J. Wawer, Vasanthi S. Viswanathan

## Abstract

Despite glutathione’s long-recognized role as a major cellular antioxidant and its central role in ferroptosis defense, inhibition of glutathione biosynthetic enzymes has received little attention as a target for the therapeutic induction of ferroptosis. Here, we report that small-molecule inhibition of glutamate–cysteine ligase (GCL), the rate-limiting enzyme of glutathione biosynthesis, selectively and potently kills cancer cells by ferroptosis. We further describe novel GCL inhibitors including KOJ-1 and KOJ-2, compounds with excellent cellular potency and pharmacological properties, representing valuable tools to study the biology of ferroptosis and glutathione.

## Introduction

Glutathione (GSH), a tripeptide first discovered in 1888, is the most abundant small-molecule thiol in animal cells and a key component of cellular antioxidant and xenobiotic defense.^1,2^ Causal or consequential perturbations of glutathione levels are a hallmark of many diseases, and upregulation of GSH is commonly observed in cancer.^3–7^ This upregulation of GSH is thought to buffer cancer cells from oncogenic reactive oxygen species (ROS) and to protect them from anti-cancer therapies. For this reason, inhibition of GSH biosynthesis has been explored sporadically over the past fifty years as a cancer vulnerability and as a means to potentiate the cell killing efficacy of cytotoxic drugs, particularly those known to involve reactive intermediates and glutathione-mediated xenobiotic detoxification.^8–13^

The discovery of ferroptosis as a coherent biological pathway has implicated a more specific role for GSH as a cytoprotective agent: that of central cofactor for the anti-ferroptotic enzyme GPX4 (Fig. 1A).^14,15^ In ferroptosis, depletion of intracellular cyst(e)ine has been shown to lead to loss of glutathione,^16,17^ which in turn compromises the function of GPX4. Without the function of GPX4, the only enzyme able to reduce complex membrane phospholipid hydroperoxides,^18–20^ cells with susceptibility to phospholipid peroxidation undergo ferroptotic death. Despite this model, as well as extensive validation of cystine transport inhibition and direct inhibition of GPX4 as mechanisms for ferroptosis induction, glutathione biosynthesis – the explicit link between cystine transport and GPX4 – has received little attention as a therapeutically relevant target for ferroptosis induction.^21–24^ In this paper, we systematically address this gap in current understanding and validate glutamate–cysteine ligase (GCL), the rate-limiting enzyme for glutathione biosynthesis, as a promising therapeutic target for ferroptosis induction in cancer.

**Figure 1:**
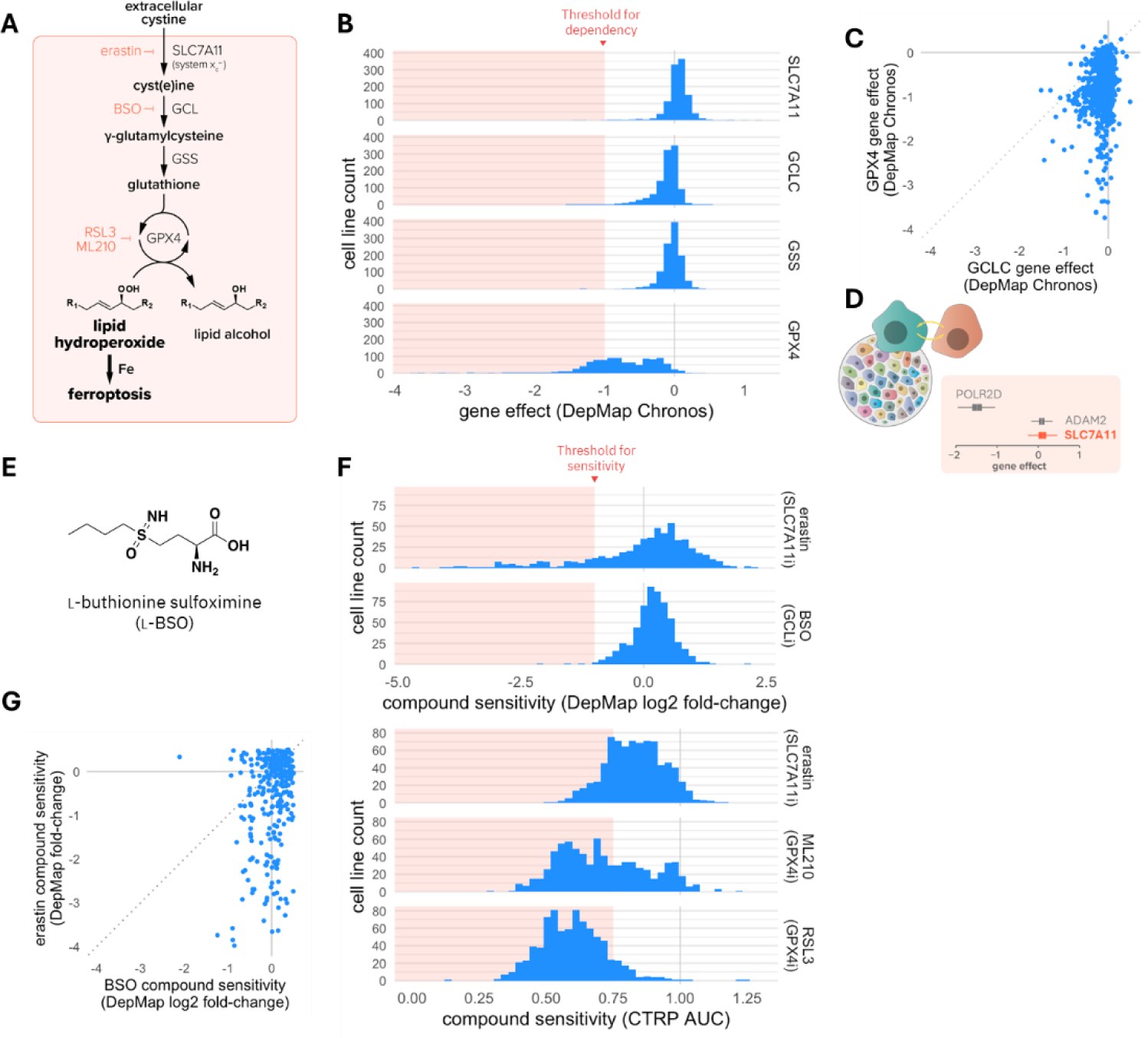
Systematic target identification approaches do not identify GCL as an anti-cancer target. (A) Schematic of the central ferroptosis pathway. GCL, glutamate-cysteine ligase; GSS, glutathione synthetase; GPX4, glutathione peroxidase 4. (B) Histogram summarizing response of 1100 cancer cell lines in DepMap to CRISPR perturbations targeting genes in the central ferroptosis pathway. (C) Scatterplot showing response of 1100 cancer cell lines in DepMap to GCLC-vs. GPX4-targeting CRISPR perturbations. (D) Schematic illustrating potential paracrine effects in pooled CRISPR screens. Colored cells represent unique gene knockouts within a pool of cells, each harboring a different gene knockout. Yellow arrows depict the exchange of metabolites between cells harboring different gene knockouts, which could account for lack of cell-autonomous dependency on certain genes such as SLC7A11. For comparison, scores for a cell-growth-essential gene (POLR2D) and non-essential gene (ADAM2) are shown. (E) Structure of L-buthionine sulfoximine (BSO), a GCL inhibiting tool compound. (F) Histogram summarizing response of 559 cancer cell lines in DepMap’s repurposing screen to BSO and erastin, a validated ferroptosis-inducing compound that inhibits the system x ^-^ cystine transporter (top). Response of 758-798 cancer cell to erastin and GPX4 inhibitors ML210 and RSL3 in DepMap’s CTRP dataset (bottom). (G) Scatterplot showing response of 559 cancer cell lines to 2.5 µM BSO vs. 10 µM erastin.

### Systematic target identification approaches do not identify GCL as an anti-cancer target

In performing a comprehensive review of the available evidence for GCL as a therapeutic target for ferroptosis, we observed that the lack of attention given to GCL was likely due to, in large part, the failure of GCL to score in important systematic cancer target identification experiments.

Now a cornerstone of public research, high-throughput target-discovery efforts such as the Broad Institute’s DepMap (www.depmap.org) have yielded powerful look-up resources that inform, genome-scale, on the impact of perturbing individual target genes in cancer models.^25–27^ DepMap is among the most comprehensive and valuable of these resources, integrating genome-scale CRISPR and shRNA perturbations, small molecule sensitivity data, and multi-omic basal characterization for over 1000 cancer cell lines from multiple sources.^25–27^ In inspecting the data for GCL in DepMap, we were struck by the fact that in this data set, while GPX4 is a strong and selective dependency across cancer cell lines, perturbation of GCLC, the catalytic subunit of GCL, is relatively inert (Fig. 1B-C). Knockout of glutathione synthetase (GSS), the enzyme required downstream of GCL for glutathione biosynthesis also does not impair cellular viability within the DepMap dataset (Fig. 1B). While this may be taken as evidence that glutathione biosynthesis enzymes are expendable for the viability of cancer cells, we observed that knockout of SLC7A11, the validated target of the ferroptosis-inducing molecule erastin and a key component of the cystine transporter system x_c_^-^, similarly does not impact cellular viability within the DepMap dataset (Fig. 1B). This raises the possibility that knockout of both SLC7A11 and glutathione biosynthesis genes are false negatives due to a technical limitation inherent in the high-throughput genetic perturbation systems required for highly multiplexed genetic screens such as those in DepMap. To illustrate, one could imagine that pathways subject to paracrine effects may fail to be uncovered by pooled CRISPR screening (Fig. 1D).^28,29^ In the case of GCLC and SLC7A11, the import of cysteine and glutathione-related metabolites from neighboring cells may circumvent the cell-autonomous loss of these proteins. Nevertheless, the failure of GCL to score as a strong cancer-cell dependency comparable to GPX4 in DepMap is likely one of the reasons why GCL has not been investigated more closely as a target for ferroptosis induction.

In addition to genetic perturbations, DepMap provides an orthogonal mechanism to identify cancer cell dependencies by making available comprehensive profiling of over 1,400 small molecules in many hundred cancer cell lines.^27^ The inclusion of the GCL inhibitor buthionine sulfoximine (BSO)^30^ (Fig. 1E) within this DepMap dataset afforded us a second opportunity to assess cancer cell dependency on GCL. Apparently consistent with genetic perturbation of GCL, pharmacological inhibition of GCL using BSO achieves little cell killing activity across the panel of 559 cancer cell lines assessed (Fig. 1F-G). However, it is important to note that the BSO dose tested in this DepMap dataset is 2.5 µM, while BSO is a poorly potent molecule often requiring high micromolar and millimolar doses in cellular assays.^24,31^ This raises the possibility that the lack of cell killing activity by BSO observed in this dataset may be due to an insufficiently high dose range. Indeed, close inspection of dose curves from a secondary DepMap screen reveals that at the top dose of 10 µM, BSO begins to show hints of activity correlating with erastin, an inhibitor of cystine transport (Supplementary Fig. 1). Nevertheless, the apparent lack of activity across most of the tested dose range and the results from the pharmacological primary screen reinforce the negative results from genetic perturbations and together fail to provide support for GCL as an anti-cancer target.

Adding to the systematic results from DepMap, many literature studies that have performed more focused interrogation of GCL as a cancer target have not uncovered significant single-agent cell-killing activity.^3,21,32–34^ Critically, most of these studies have been conducted in epithelial-state, ferroptosis-insensitive cancer cells, which do not have the capacity to die via ferroptosis. It is therefore possible that performing such studies in mesenchymal-state cancer cells could yet reveal the capacity of GCL inhibition to trigger ferroptotic death. Furthermore, the few recent studies that have uncovered single-agent cell killing activity by BSO, usually limited to specific contexts, do not establish a link to ferroptosis, and thus do not identify the scope and rationale for GCL inhibition in cancer that is informed by the current understanding of ferroptosis susceptibility across cancers.^23,35,36^

As we sought to investigate GCL as a potential ferroptosis target, we took care to learn from the historical results described above and to account for potential technical artifacts and confounders that may have impeded previous efforts. Specifically, we sought to perform experiments in cells known to be susceptible to ferroptosis. For pharmacological studies, we ensured the use of a dose range appropriately optimized for the tool compound BSO. For genetic perturbations, we prioritized arrayed rather than pooled experimental designs to avoid artifacts (including potential paracrine effects) arising in mixed populations.

### BSO leads to robust ferroptosis induction

To test whether GCL inhibition can lead to ferroptosis, we performed dose-titration experiments using a wide range of concentrations of BSO in ferroptosis-susceptible (Fig. 2A) and ferroptosis-insensitive cell lines (Fig. 2B). In the ferroptosis-susceptible cell lines, BSO led to strong cell-killing effects, as assessed by both CellTiter-Glo measurements and cell imaging (Fig. 2A, 2C). Importantly, in two out of three of these sensitive cell lines, the concentration of BSO required for cell-killing activity was well beyond the 10 µM top dose used in DepMap for BSO, providing one potential explanation for the lack of activity by BSO seen in DepMap. Meanwhile, BSO across the full dose range had no impact on the viability of ferroptosis-insensitive cells (Fig. 2B).

**Figure 2:**
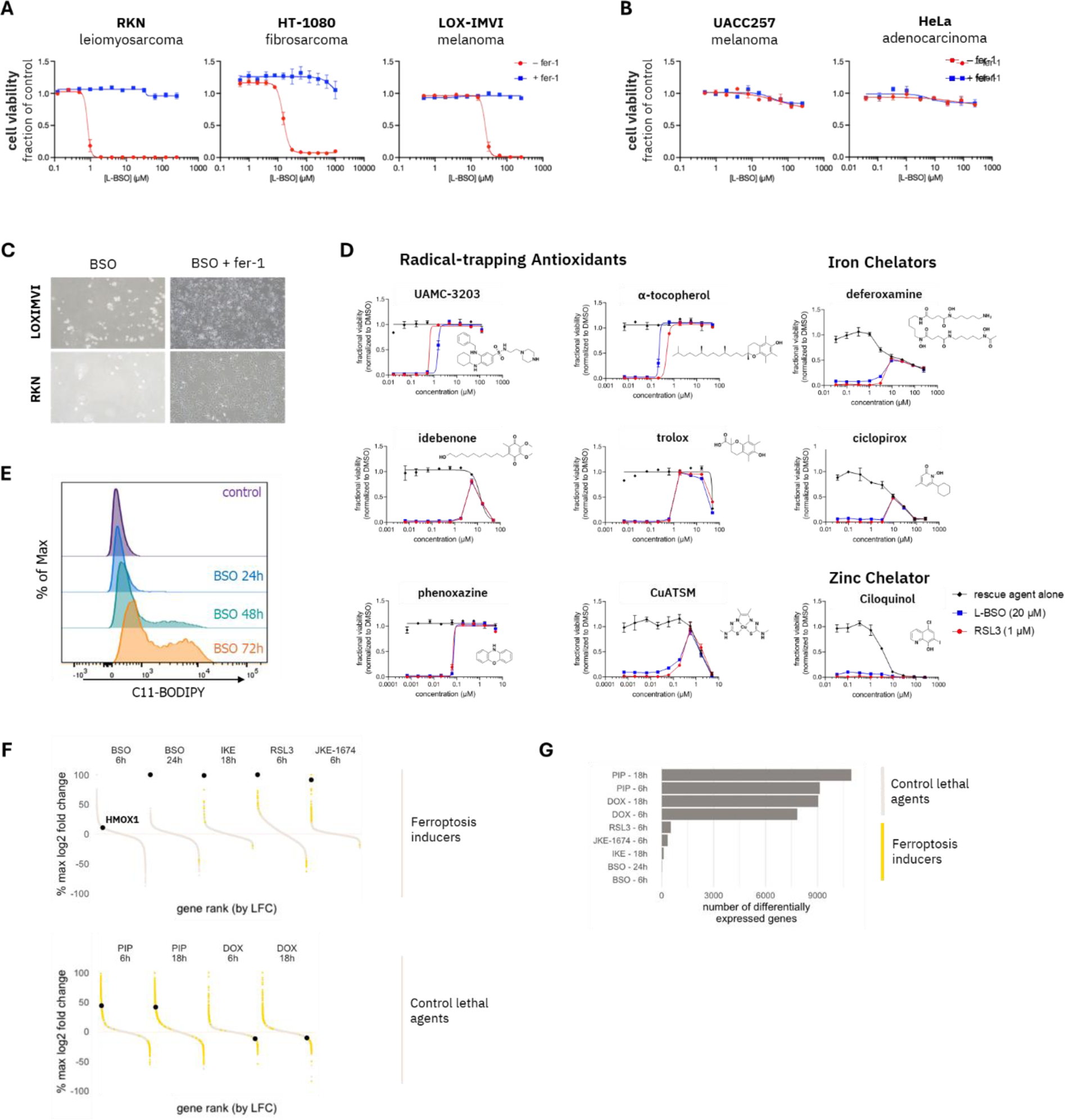
BSO induces robust ferroptosis induction. Dose-response curves showing cell-killing by BSO in (A) ferroptosis-sensitive cell lines and (B) ferroptosis-insensitive cell lines. Red curves reflect treatment with BSO alone; blue curves reflect co-treatment with BSO and 1.5 µM ferrostatin-1. Data are plotted as mean ± s.d. for n = 2 replicates; 72 h incubation time. (C) Brightfield microscopy images showing cell killing by 100 µM BSO in two ferroptosis sensitive cell lines and rescue of BSO-mediated cell killing by 1.5 µM ferrostatin-1 co-treatment. (D) Dose-response curves in RKN cells demonstrating the capacity of an extended panel of radical-trapping antioxidant and iron chelators to rescue from ferroptosis induced by either BSO or the GPX4 inhibitor RSL3. Experiments performed in RKN cells with 72-hour incubation time. (E) Time-dependent induction of lipid peroxidation in RKN cells treated with 100 µM BSO, as assessed using C11-Bodipy via flow cytometry. (F) Gene expression changes in RKN cells treated with a panel of ferroptosis-inducing compounds and control lethal agents for the indicated durations. Each dot represents an individual transcript. Dots colored in yellow are significantly altered relative to DMSO control, BH-adjusted p < 0.01 Induction of HMOX1 is a ferroptosis-selective transcriptomic response in these cells. BSO, 10 µM; IKE, imidazole ketone erastin, 1µM; RSL3, GPX4 inhibitor, 0.1 µM; JKE-1674, GPX4 inhibitor, 1 µM; PIP, piperlongumine, 10 µM; DOX, doxorubicin, 1 µM. (G) Bar plot showing the number of genes differentially expressed relative to DMSO in each treatment condition from the experiment shown in (F).

Viability effects of BSO were rescued by ferrostatin-1, a highly specific and well-validated ferroptosis preventing agent,^16,37^ thus providing initial evidence for the ferroptotic nature of BSO-induced cell death (Fig. 2A-C). We confirmed this via co-treatment with an extensive panel of ferroptosis-preventing agents,^16,38–41^ which rescued BSO in a manner identical to the gold-standard ferroptosis inducer RSL3 (GPX4 inhibitor)^14^ (Fig. 2D). Time-dependent induction of lipid peroxidation in cells treated with BSO provided further confirmation of ferroptotic cell death (Fig. 2E). Cells treated with BSO also exhibited a transcriptomic signature consistent with other ferroptosis-inducing perturbations. To demonstrate this, we harvested cells dosed with diverse ferroptosis-inducing and control lethal agents at comparable time points in their transition to cell death, as monitored by microscopy. Ferroptosis inducers, including BSO, induced remarkably few transcriptional changes as compared to control lethal agents (Fig. 2F-G, Supplementary Fig. 2A). However, all ferroptosis inducers, including BSO, led to characteristic induction of HMOX1, a known ferroptosis response gene (Fig. 2F, Supplementary Fig. 2A).

### Ferroptosis induction by BSO is mediated via GCL inhibition

Having established BSO as an inducer of ferroptotic cell death, we sought to confirm GCL inhibition as the mediating mechanism for the effects of BSO. BSO and other sulfoximine analogs are mechanism-based inhibitors of GCL.^2,30^ Upon binding, the sulfoximine nitrogen of these compounds is phosphorylated by the GCL enzyme, mimicking the transition state of the ligation reaction between cysteine and glutamate.^30,45^ The resulting phosphorylated sulfoximine is tightly bound to the enzyme with a slow off rate, affording BSO and analogs with remarkable selectivity for GCL over the mechanistically related enzymes glutathione synthetase (Supplementary Fig. 3) and glutamine synthetase.^46^ However, even mechanistically specific molecules can impact cells primarily via off-target effects when used at the high concentrations required for ferroptosis induction by BSO. To probe this possibility, we aimed to establish close alignment between the dose-response of BSO for GCL binding, glutathione depletion, and ferroptosis induction, hypothesizing that any significant discrepancy would indicate the involvement of non-GCL off-target effects of BSO as responsible for ferroptosis induction. We established a cellular thermal shift assay (CETSA)^47^ for both the catalytic and modifier subunits of GCL, hereafter GCLC and GCLM, respectively, to report on direct target engagement by BSO (Fig. 3A) and validated methods for measurement of cellular glutathione, a downstream product of GCL (Fig. 3B). Using these assays, we performed dose-titration experiments with BSO and compared these with the dose-titration curves for ferroptosis induction by BSO in the same cell line (Fig. 3C). The results reveal a remarkable concordance between GCL target engagement, enzyme inhibition and ferroptosis induction, consistent with GCL inhibition as the mechanism by which BSO induces ferroptosis. Furthermore, glutathione depletion precedes ferroptosis induction, supporting its role as a cause rather than a consequence of oxidative cell death (Fig. 3D). A broader panel of glutathione-related metabolites,^48^ including γ-glutamylcysteine,^48^ the immediate product GCL, measured by mass spectrometry confirmed the robust and specific effects of GCL inhibition (Fig. 3E).

**Figure 3.**
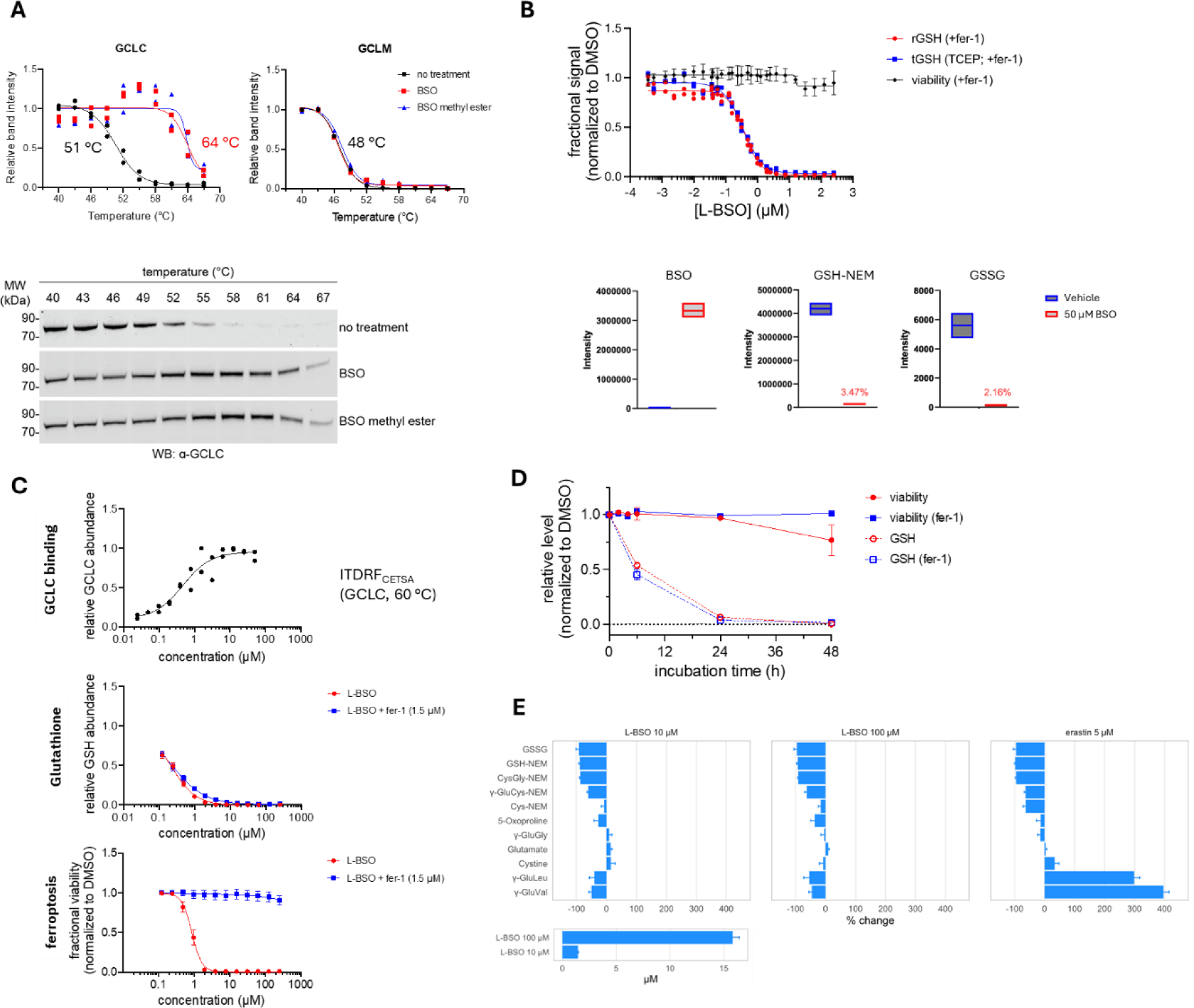
Ferroptosis induction by BSO is mediated via GCL inhibition. (A) Western blot and quantitation of protein levels showing cellular melting curves (CETSA) for the catalytic (GCLC) and modifier (GCLM) subunits of GCL in the presence or absence of 500 µM BSO or BSO methyl ester after a 4 h incubation. Data plotted are individual replicates from 2 independent experiments. (B) Kit-based (top) and mass-spec based (bottom) measurement of reduced glutathione (rGSH) and total glutathione (tGSH) in cells treated for 48 **h** with varying concentrations of BSO in the presence of 1.5 µM fer-1 to maintain cellular viability. Data plotted are two replicates. (C) Cellular GCLC stabilization (CETSA), cellular glutathione levels and ferroptosis induction as a function of BSO dose. CETSA data plot an individual replicate of representative data following 3 **h** incubation with compound. Glutathione measurements and viability measurements were performed following 72 **h** of treatment with BSO. Data plotted are mean ± s.d.; n = 16 for glutathione measurements and n = 22 for viability measurements. (D) Time-course experiment measuring cellular glutathione levels vs. induction of ferroptosis in response to 10 µM BSO over a period of 48 hours. (E) Confirmation by mass spectrometry of changes in glutathione levels as well as other GCL-specific metabolomic changes in cells treated with the indicated concentration of BSO for 48 hours. (A) was performed in KP4 cells. All other experiments were performed in RKN cells.

### Novel BSO analogs preserve tight correspondence between GCL inhibition and cellular ferroptosis induction

Structure–activity relationship (SAR) studies are an orthogonal means to gain insight into the mechanism of action of molecules. Reported SAR studies of BSO to date have been limited in scope. To facilitate more in-depth SAR studies for BSO analogs that would probe the link between GCL inhibitory activity and cellular ferroptosis induction, we established a biochemical enzyme activity assay using recombinant GCL protein (Fig. 4A). We next synthesized a series of BSO analogs probing SAR of the butyl chain, amino acid, and sulfoximine portions of the molecule (Fig. 4B). Monitoring enzyme inhibition confirmed that the branching pattern and presence of a hydrophobic anchoring group (e.g., trifluoromethyl) within the cysteine thiol binding pocket are important for GCL inhibitory activity. Treating ferroptosis-susceptible cancer cells with this panel of BSO analogs revealed that GCL inhibitory activity and ferroptosis induction are tightly correlated (Fig. 4B). Most modifications to the amino acid portion of BSO were not tolerated, including backbone homologation that incorrectly positions the sulfoximine group for phosphorylation by the enzyme and results in a biochemically inert analog unable to induce ferroptosis. Furthermore, assessment of all four BSO isomers^49^ revealed concordance in potency of purified GCL inhibition, cellular glutathione depletion, and ferroptosis induction (Fig. 4C). These SAR results provide compelling evidence that GCL inhibition is the mechanism by which BSO induces ferroptosis.

**Figure 4.**
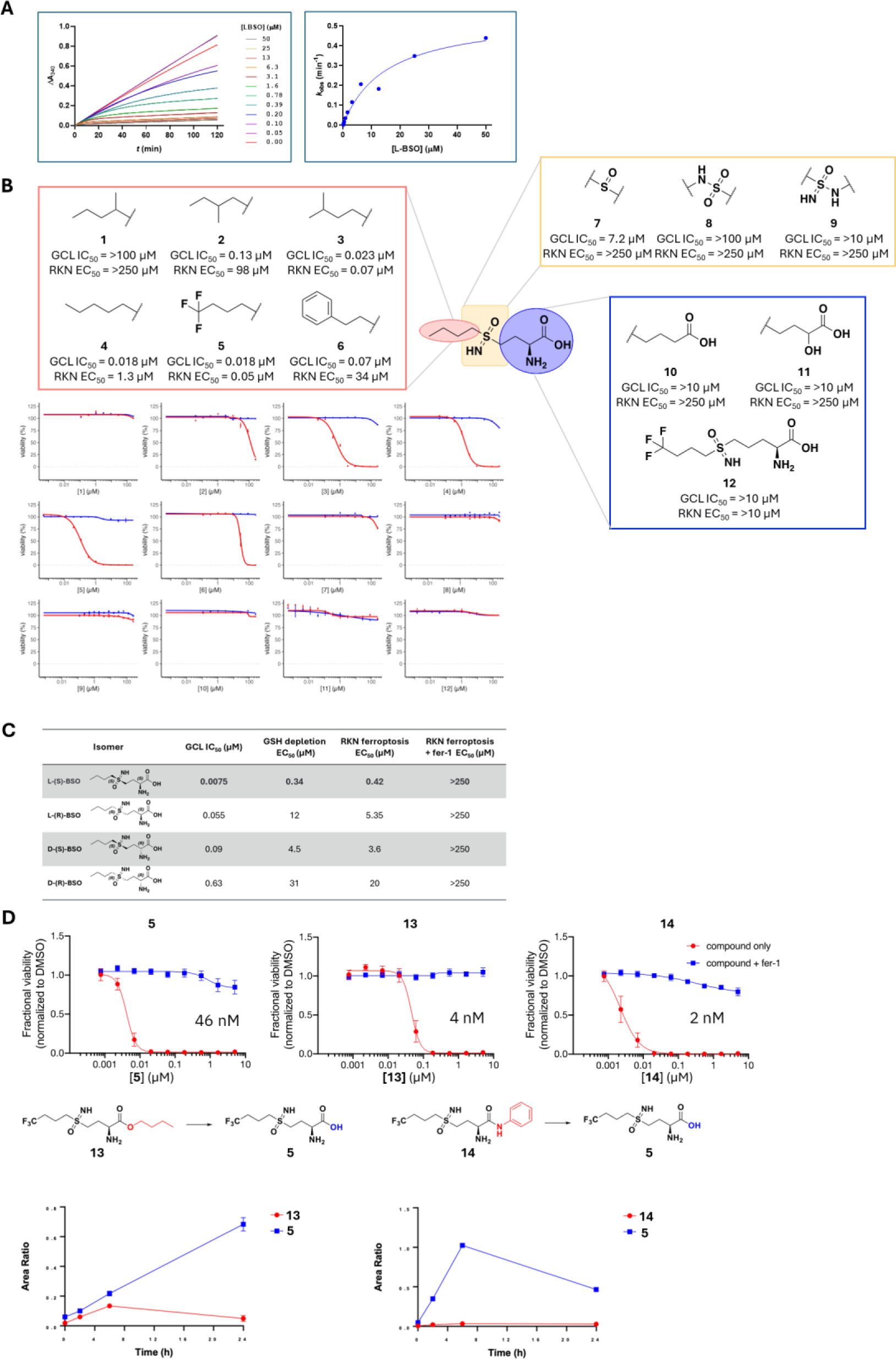
BSO analogs preserve tight correspondence between GCL inhibition and cellular ferroptosis induction. (A) Time-dependent inactivation of human GCL by BSO. (B) Summary of BSO structure-activity relationship (SAR) studies measuring inhibition of purified GCL enzyme activity and ferroptosis induction in RKN cells, in the presence or absence of 1.5 µM ferrostatin-1. (C) Stereochemical SAR of BSO supports correlation between inhibition of purified GCL enzymatic activity, cellular glutathione depletion and ferroptosis induction. (D) Prodrug analogs get converted into the active species in cells and induce ferroptosis with greater potency.

Our SAR studies also revealed a strategy for increasing cellular exposure to sulfoximine GCL inhibitors, which would improve ferroptosis induction potency. The highly polar BSO and related sulfoximines have low membrane permeability and most likely require active transport for efficient cellular accumulation. We hypothesized that prodrug analogs of sulfoximine GCL inhibitors would have improved membrane permeability and could achieve higher intracellular concentrations more quickly, which should translate into increased potency for ferroptosis induction. To test this, we generated prodrugs by converting the carboxylic acid group into esters and amides, modifications that diminish the ability of these analogs to inhibit purified GCL enzyme (Fig. 4D). However, in cells, the prodrugs were rapidly converted into the active free-acid form and achieved more potent depletion of cellular glutathione and ferroptosis induction (Fig. 4D).

Emerging from these efforts is KOJ-1, a potent cellular GCL inhibitor with improved pharmacological properties that represents an excellent tool compound for interrogating ferroptosis and glutathione biology *in vitro*. It is important to note that ester prodrugs like KOJ-1 are not appropriate for inhibiting purified GCL biochemically. *In vivo*, hydrolysis of KOJ-1 occurs rapidly, which negates the potency advantage the ester group provides for ferroptosis induction in cells. To address this limitation, we developed KOJ-2, a non-prodrug which, compared to BSO, exhibits improved biochemical inhibition of GCL, increased ferroptosis induction potency in cells, and better systemic exposure in mice following oral administration (Supplementary Figure 4). Preliminary observations support the *in vivo* bioavailability and bioactivity of KOJ-2 (Supplementary Figure 4) and provide a rationale for further chemical optimization and *in vivo* exploratory studies. KOJ-2 was independently reported in a recent patent and similar enzyme inhibition and *in vivo* bioactivity to what we report was observed.^50^

### Genetic perturbation of GCL induces ferroptosis

While our emphasis has been on pharmacological perturbation of GCL and chemical-biology approaches to validate the target, genetic approaches also confirm that loss of GCL can trigger ferroptotic cell death. To systematically perturb nodes in the central ferroptosis pathway (Fig. 1A) genetically, we employed direct ribonucleoprotein (RNP) mediated delivery of gene-targeting CRISPR-Cas9 complexes into two ferroptosis-susceptible cell lines and one ferroptosis-insensitive cell line. Loss of GCL resulted in ferrostatin-1-rescuable cell killing in ferroptosis-susceptible cell lines, in a manner analogous to loss of GPX4 (Fig. 5A). Interestingly, loss of SLC7A11 as well as loss of glutathione synthetase (GSS), the enzyme downstream of GCL that is required for glutathione biosynthesis, each also resulted in ferroptosis induction. These results stand in contrast to the knockout phenotypes for these genes seen in DepMap (Fig. 1B), suggesting that there may indeed be technical aspects of the highly multiplexed pooled CRISPR-screening approach employed by DepMap (e.g., paracrine effects) that lead to systematic false negatives for certain classes of ferroptosis targets. Experiments using inducible shRNAs targeting GCLC yielded similar results (Fig. 5B) to those seen with GCLC-target CRISPR RNPs (Fig. 5A). In contrast, CRISPR RNP particles targeting GCLM, the modifier subunit of GCL which is not required for glutathione biosynthesis, did not lead to decreased viability (Fig. 5A).

**Figure 5.**
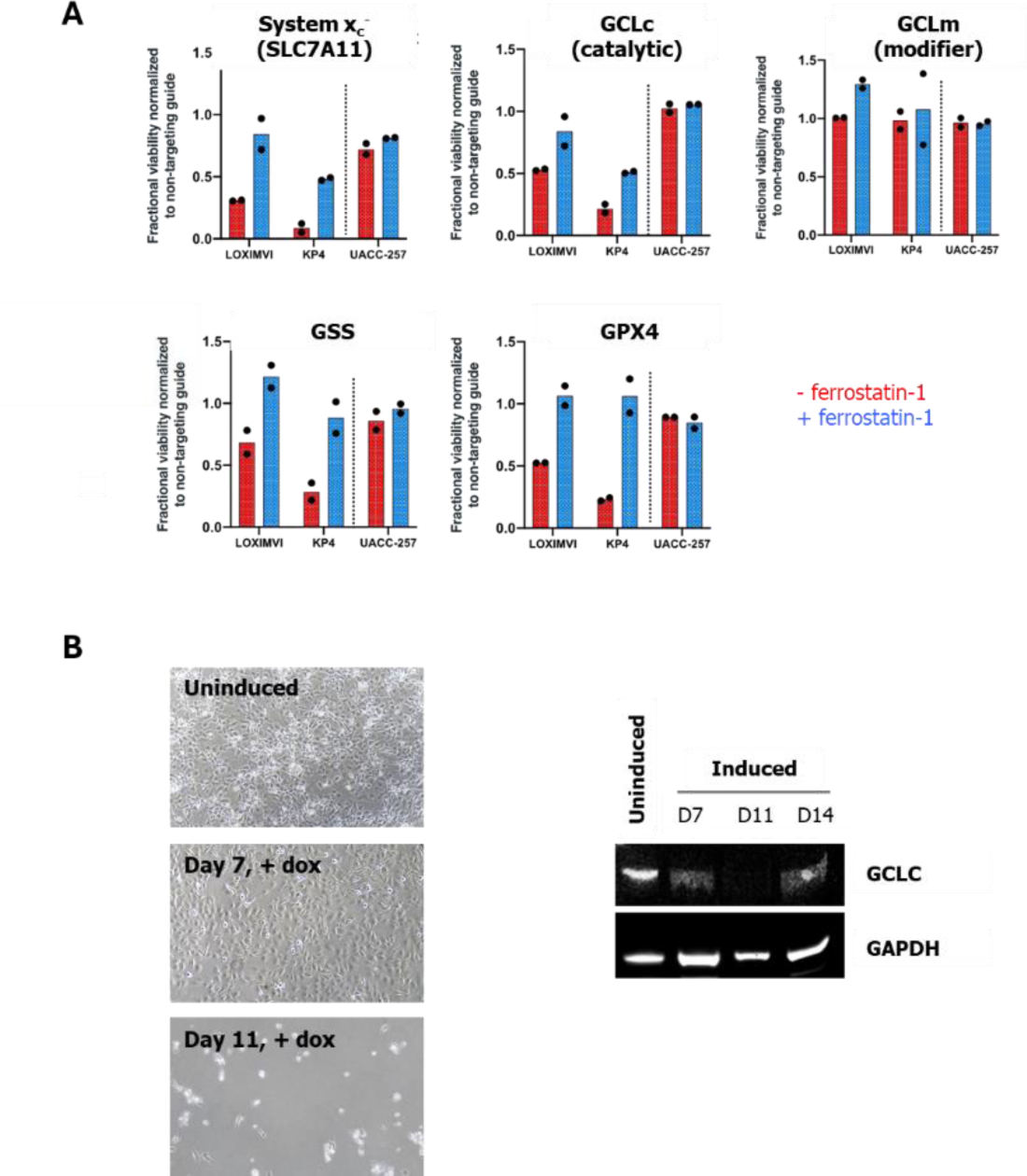
Genetic perturbation of GCL induces ferroptosis. (A) CRISPR-RNP based knockout of target genes in the central pathway of ferroptosis, including the catalytic subunit of GCL (GCLC), induce ferroptosis in ferroptosis-sensitive cells (LOXIMVI, KP4), while having no impact on ferroptosis-insensitive cells (UACC-257). Data plotted are two technical replicates, 7 days post-transfection. (B) Brightfield microscopy images demonstrating cell death upon induction with doxycycline of RKN cells transduced with a doxycycline-inducible GCLC-targeting shRNA.

### GCL inhibition induces ferroptotic cell death across a large panel of cancer cell lines

Having established that GCL inhibition can induce ferroptotic cell death, we next sought to understand how relevant this mechanism might be across cancer cell lines more broadly. To address this, we tested a pro-drug analog of BSO across an initial panel of 35 cancer cell lines (Fig. 6A). Based on metrics of absolute sensitivity to GCL inhibition as well as rescue by ferrostatin-1 cotreatment, we find that about one third of the cancer cell lines tested in this panel undergo ferroptosis in response to GCL inhibition (Fig. 6A).

**Figure 6.**
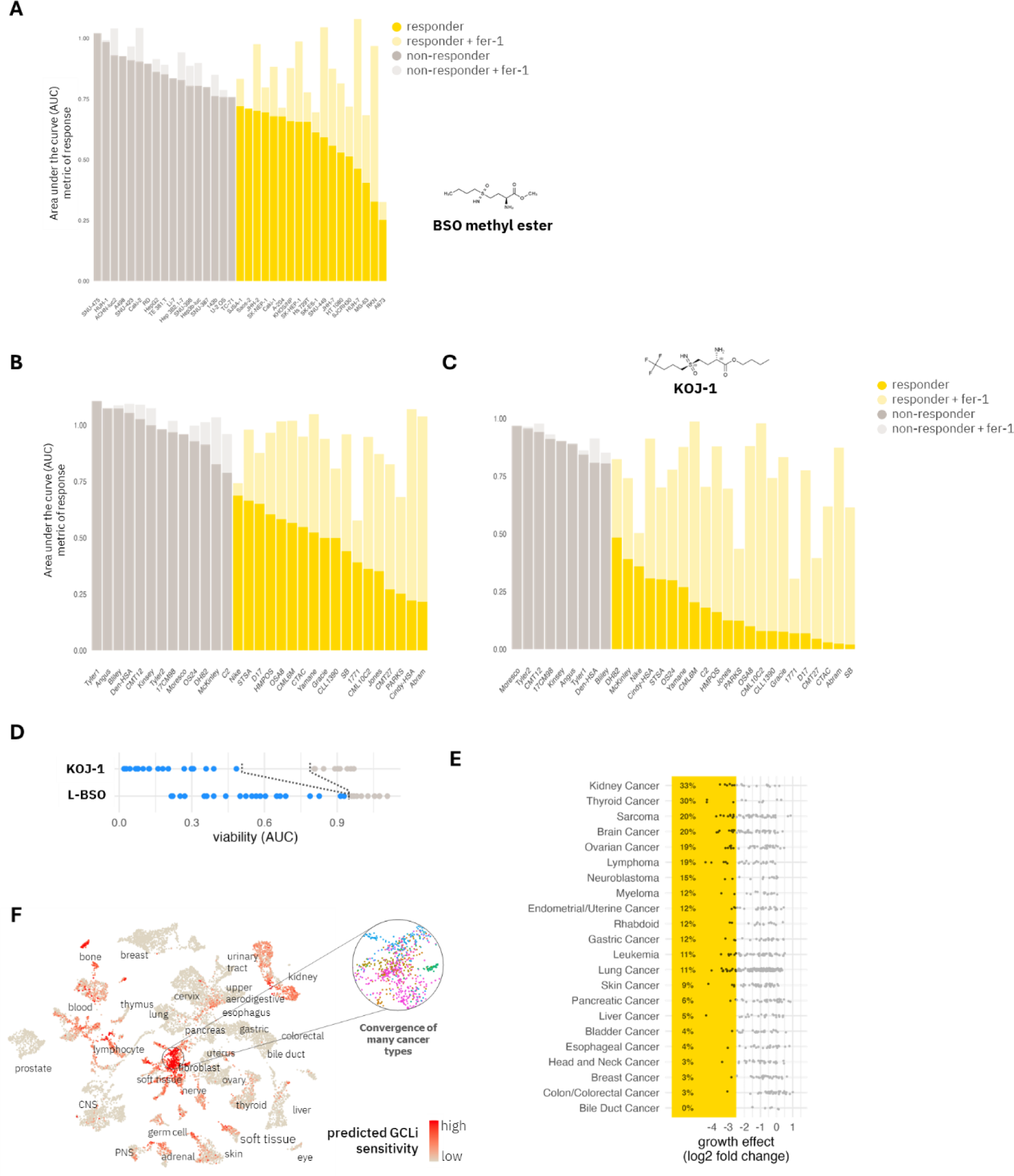
GCL inhibition induces ferroptotic cell death across a large panel of cancer cell lines. (A) Bar graph summarizing responses of 35 human cancer cell lines to BSO methyl ester. Data plotted are an AUC metric of sensitivity integrated across 9 dose points ranging from 38 µM to 250 µM. Darker bars in the foreground reflect treatment with BSO methyl ester alone. Lighter bars in background reflect cotreatment with BSO methyl ester and 1.5 µM ferrostatin-1. Cell lines highlighted in yellow are deemed sensitive to BSO methyl ester using an AUC cut-off of 0.75. (B) Bar graph summarizing response of 31 canine cancer cell lines to BSO and (C) to KOJ-1, a more potent BSO analog. Details of visualizations in (B) and (C) are same as for (A). (D) KOJ-1 expands the selectivity window for ferroptosis induction (blue dots) compared to BSO. (E) Summary of cancer cell line responses by cancer type to 50 µM BSO across Broad Institute PRISM 723 cancer cell line panel. Cancer types with more than 5 cell lines are shown. (F) UMAP plot, based on Celligner tool, depicting predicted sensitivity of 12,236 patient tumors and 1249 cell lines to GCL inhibition. Inset highlights a region of heightened sensitivity to ferroptosis that reflects a highly conserved undifferentiated cancer cell state that arises in many different cancer types.

We recently reported the validation of canine cancer cell lines as translationally relevant models for investigation of ferroptosis induction.^51^ The canine cancer context is especially significant for the study of ferroptosis given that mesenchymal and undifferentiated cancer types, which are known to have heightened sensitivity to ferroptosis, are more prevalent in canine oncology than in the human setting.^52,53^ Testing BSO across a panel of 31 available canine cancer cell lines^54^ revealed a striking rate of ferroptotic response to GCL inhibition, with mesenchymal cancer cell lines showing greatest sensitivity, consistent with current understanding of ferroptosis susceptibility across cancer types (Fig. 6B). A more potent analog of BSO induces ferroptosis in the most sensitive canine cancer cell lines with single-digit nanomolar potency (Fig. 6C) and significantly expands the selectivity window for ferroptosis induction between canine cancer cell lines in a mesenchymal cell state vs. those with epithelial characteristics (Fig. 6D, Supplementary Fig. 5).

To probe cancer cell line vulnerability to GCL inhibition across a much broader spectrum of cancer cell lines, we profiled the response of the Broad Institute’s PRISM 723 cancer-cell-line panel^27^ to 50 µM BSO, a five-fold higher concentration than the 10 µM top dose used for BSO in the publicly available DepMap dataset. Our results demonstrate that increasing the dose of BSO is sufficient to uncover considerable growth-inhibitory effects across cell lines representing many different cancer types (Fig. 6E). The well-known hallmarks of ferroptosis susceptibility,^55^ including heightened sensitivity of sarcomas and kidney cancers and relative insensitivity of highly epithelial tumor types, are recapitulated in this data set.

Subsets of other major cancer types with the potential to be molecularly defined also demonstrate meaningful sensitivity to GCL inhibition. Moreover, as seen in the canine cancer cell line panel (Fig. 6B-C), the use of a more potent drug-like GCL inhibitor, while retaining the overall pattern of sensitivity uncovered with 50 µM BSO, would be expected to dramatically improve upon the depth and selectivity of responses to BSO seen here. To investigate how patterns of cancer cell line sensitivity to GCL inhibition may inform sensitivity of patient tumor types, we trained a computational model to predict sensitivity to GCL inhibition from gene-expression values. To harmonize expression values between cell lines and patient samples and visualize results, we made use of Celligner,^56^ a tool recently developed for this purpose. The resulting UMAP plot highlights broad indication scope for GCL inhibition in cancer based on the *in vitro* data we have reported in this paper (Fig. 6F) and reinforces the heightened ferroptosis sensitivity of a highly undifferentiated cell state that cancers arising from many lineages converge upon (Fig. 6F inset).^55,57^

## Discussion

The results reported herein comprise a comprehensive *in vitro* validation of GCL as a target for ferroptosis induction in cancer. More broadly, our findings eliminate a long-standing enigma within the central ferroptosis pathway, namely that upstream cyst(e)ine depletion and downstream GPX4 inhibition readily induce ferroptosis, yet inhibition of glutathione biosynthesis – the direct link between cyst(e)ine and GPX4 – appears to be ineffective. By demonstrating the capacity of GCL inhibition to induce ferroptosis robustly and extensively, our findings provide a long-sought pillar further cementing the current model of the central Cys-GSH-GPX4 ferroptosis pathway.

While the GCL inhibitor BSO has poor pharmacokinetic properties that preclude it from being a drug-like molecule,^13^ human clinical trials in prior decades demonstrated tolerability and glutathione-depleting activity of BSO administered via continuous IV infusion.^8^ These observations motivated our efforts to further optimize this chemotype and deliver much-improved and cellularly validated GCL-inhibiting tool compounds for the first time in the 50-year history of this target. These novel probe compounds, along with the assays and technical parameters for studying GCL inhibition that have been optimized and reported as part of the present work, provide the scientific community with important new tools to carefully interrogate GCL biology and its therapeutic applications.

## Author contributions

J.K.E, M.J.W., and V.S.V. conceived of and designed research; P.C., Y.Y.S., L.L.C., K.D., R.A.V., E.P. supported cell culture maintenance and small molecule sensitivity profiling; P.C. performed CRISPR and shRNA experiments; L.L.C. performed CETSA experiments; L.F. and J.K.E. conceived of SAR studies; L.F. and J.H.J directed chemical synthesis and mouse PK experiments; G.T., S.B., and A.P. implemented enzyme inhibition assays; Y.Y.S. performed flow cytometry analyses; G.X. performed mass-spec-based metabolite measurements; J.N. analyzed RNA-seq data; J.K.E., M.J.W. and V.S.V. analyzed data and wrote the paper.

## Acknowledgements

We thank Kay Ahn for helpful conversations.

## Competing Interest Statement

V.S.V is a co-founder and equity holder of Kojin Therapeutics. All authors are equity holders of Kojin Therapeutics and are either current or past employees of Kojin Therapeutics. J.K.E, L.F., J.H.J. and V.S.V. are inventors of patents related to ferroptosis and GCL.

---

**Supplementary Figure 1.**
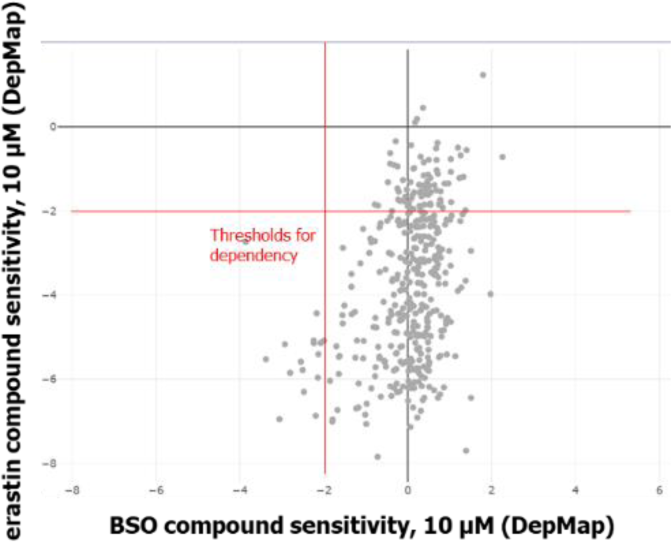
Scatterplot showing response of 1100 cancer cell lines in DepMap to GCLC-vs. GPX4-targeting CRISPR perturbations.

**Supplementary Figure 2.**
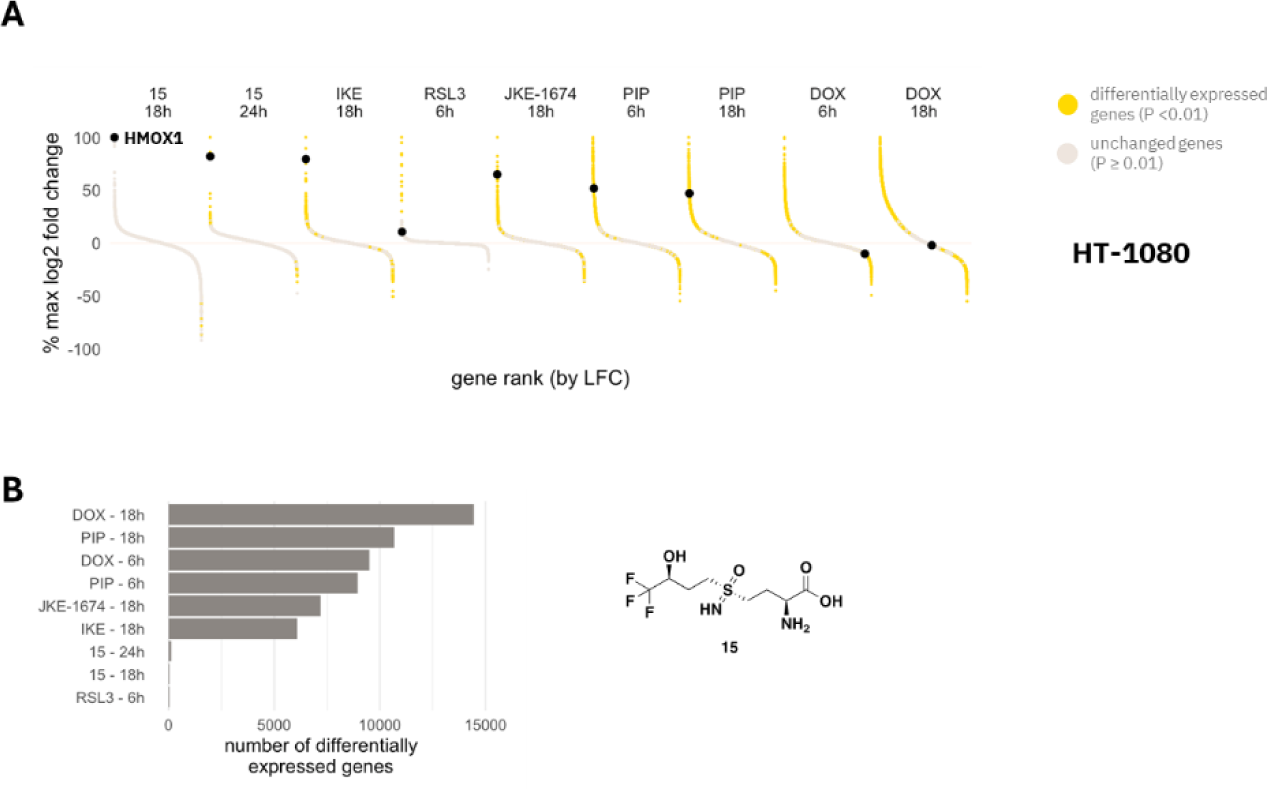
Transcriptomic changes in HT-1080 cells in response to treatment with a panel of ferroptosis-inducing and control compounds. Gene expression changes in HT1080 cells treated with a panel of ferroptosis-inducing compounds and control lethal agents for the indicated durations. Each dot represents an individual transcript. Dots colored in yellow are significantly altered relative to DMSO control, BH-adjusted p < 0.01. Induction of HMOX1 is a ferroptosis-selective transcriptomic response in these cells. BSO analog, 1 µM; IKE, imidazole ketone erastin, 1µM; RSL3, GPX4 inhibitor, 0.1 µM; JKE-1674, GPX4 inhibitor, 1 µM; PIP, piperlongumine, 10 µM; DOX, doxorubicin, 1 µM. Bar plot shows the number of genes differentially expressed relative to DMSO in each treatment condition.

**Supplementary Figure 3.**
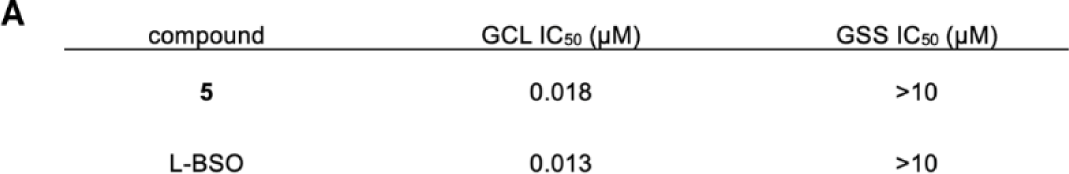
Scatterplot showing inhibitory activity of compounds against GCL vs. GSS.

**Supplementary Figure 4.**
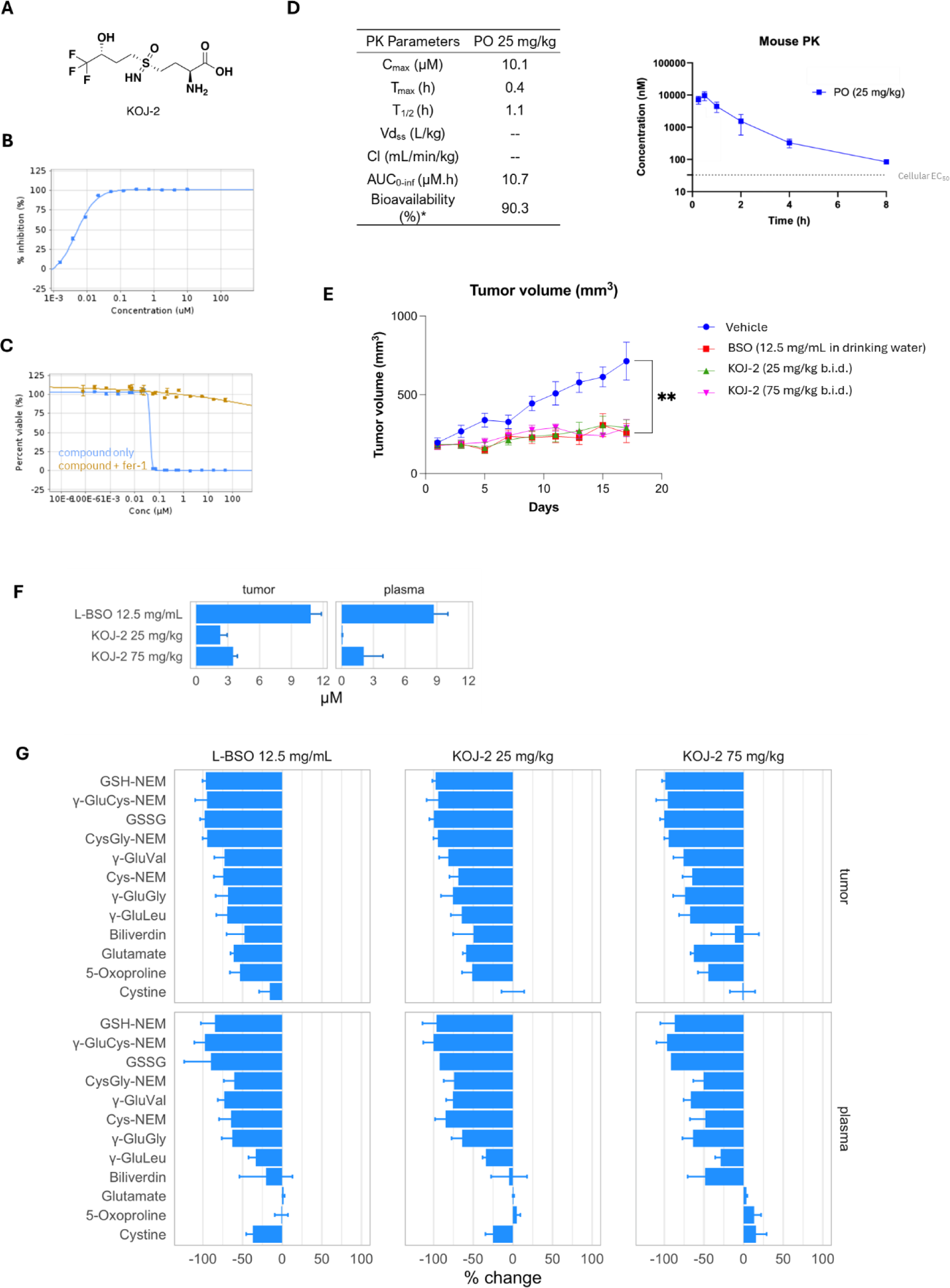
KOJ-2 is an improved in vivo tool compound. (A) Structure of KOJ-2. (B) Inhibition of purified GCL by KOJ-2. (C) Dose response data showing induction of ferroptosis by KOJ-2 in RKN cells. (D) Mouse PK characterization of KOJ-2. (E) Preliminary observations in HT1080 xenograft-bearing nude mice (n = 10) demonstrating (F) bioavailability and (G) bioactivity of orally administered KOJ-2 comparable to BSO administered in drinking water.

**Supplementary Figure 5.**
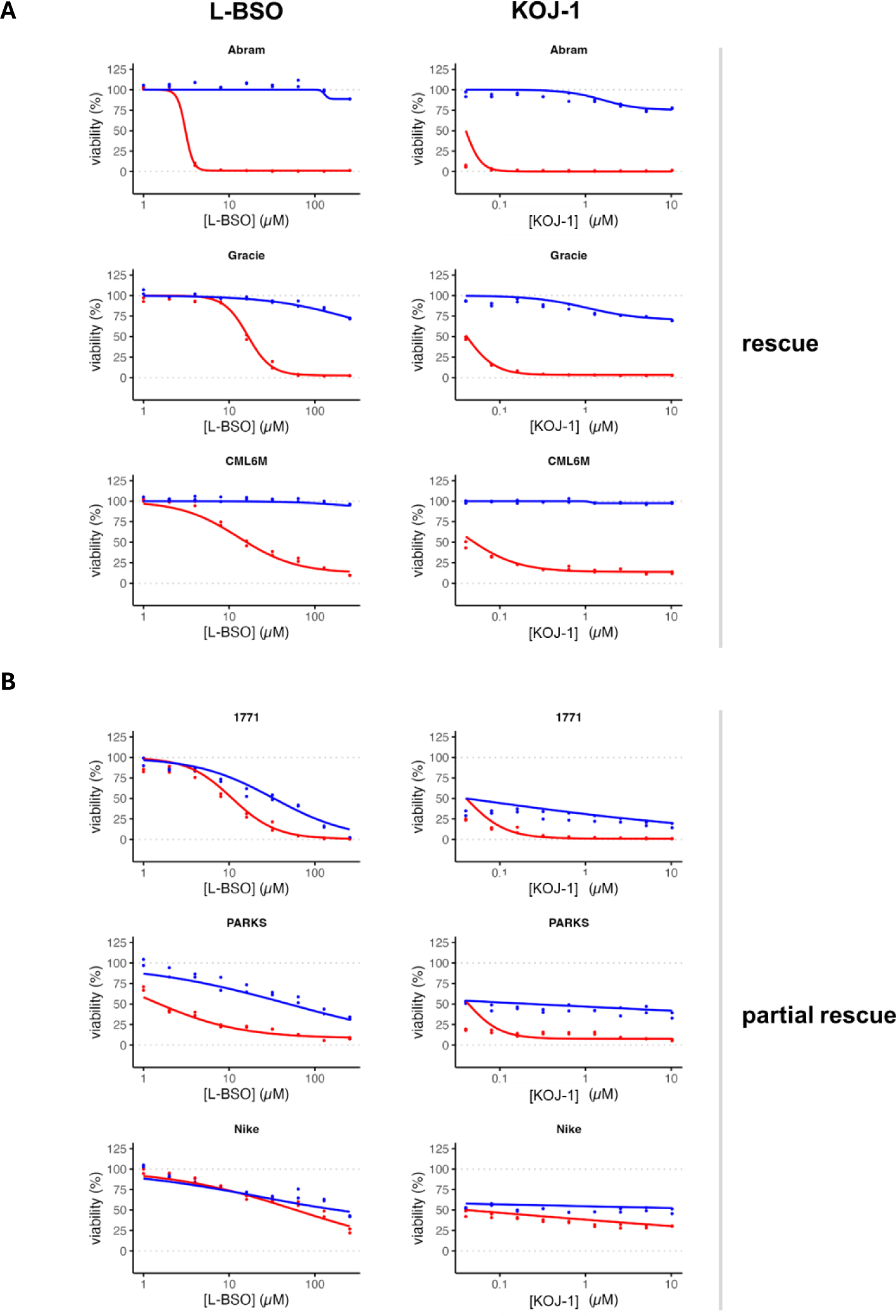
Dose-response curves illustrating (A) examples of canine cancer cell lines in which cell-killing effects of GCL inhibition are fully rescued by ferrostatin-1 vs. (B) canine cancer cell lines in which ferrostatin-1 prevents cell killing due to GCL inhibition but does not fully rescue growth proliferation (i.e., partial rescue). Red curves reflect treatment with GCL inhibitor alone; blue curves reflect co-treatment with GCL inhibitor and 1.5 µM ferrostatin-1. Data are plotted as two individual replicates; 72 h incubation time.

